# Soil, ocean, hot spring, and host-associated environments reveal unique selection pressures on genomic features of bacteria in microbial communities

**DOI:** 10.1101/2021.04.05.438506

**Authors:** Peter F. Chuckran, Bruce Hungate, Egbert Schwartz, Paul Dijkstra

## Abstract

Free-living bacteria in nutrient limited environments often exhibit small genomes which curb the cost of reproduction – a phenomenon known as genomic streamlining. Streamlining has been associated with a suite of traits such as reduced GC content, fewer 16S rRNA copies, and a lower abundance of regulatory genes, such as sigma (σ) factors. Here, we analyzed these traits from 116 publicly available metagenomes derived from marine, soil, host associated, and thermophilic communities. In marine and thermophilic communities, genome size and GC content declined in parallel, but GC content was higher in thermophilic communities. In soils, the relationship between genome size and GC content was negative, suggesting a different selection pressure on genome size and GC content in soil bacteria. The abundance of σ-factors varied with average genome size, ecosystem type, and the specific functions regulated by the sigma factor. In marine environments, housekeeping and heat-shock σ-factor genes (*rpoD* and *rpoH* respectively) increased as genome size declined, and σ-factor responsible for flagella biosynthesis (*fliA*) decreased, suggesting a trade-off between nutrient conservation and chemotaxis. In soils, a high abundance of *fliA* and the stress response σ-factor gene (*rpoS*) was associated with smaller average genome size and often located in harsh and/or carbon-limited environments such as deserts or agricultural fields – suggesting an increased capacity for stress response and mobility in nutrient-poor soils. This work showcases how ecosystem-specific environmental constraints force trade-offs which are then embedded in the genomic features of bacteria in microbial communities, specifically genome size, GC content, and regulatory genes, and further highlights the importance of considering these features in microbial community analysis.

## BACKGROUND

Assessing microbial communities through a trait-based framework highlights important relationships between microbes and their environment which may not be detectable through taxonomic analyses alone [1–6]. Notably, genomic characteristics such as genome size, GC content, number of regulatory genes, and number of 16S rRNA gene copies, have been shown to be indicators for growth rates [7], life history strategies [8] and population dynamics [9] of bacteria. Relationships between genomic features and environmental factors such as nutrient usage [9–11], aboveground cover [12, 13], temperature [14], and precipitation [15] have demonstrated the potential utility of genomic traits for assessing the relationship between bacteria and their environment.

The genome size of free-living bacteria may be reduced by a process called genomic streamlining, wherein nutrient limitation selects for smaller genomes as a way to reduce the cost of reproduction [16]. Streamlined genomes are associated with a number of traits which also reduce reproductive costs, most notably a lower GC content, fewer regulatory genes (specifically those encoding σ-factors), smaller intergenic spacer regions, and fewer 16S rRNA gene copies [11]. Consequently, bacteria with streamlined genomes tend to have a higher resource use efficiency and lower growth rates compared to bacteria with larger genomes and more rRNA gene copies [17]. Although experimental genome reduction has shown mixed relationships between genome size and growth rate [18, 19], many studies have found evidence supporting this relationship [7, 20–22]. In this way, genome size could be a functionally useful trait in predicting ecological phenomena such as growth rate. The prevalence of small genomes has long been recognized in marine systems [23] where the streamlined SAR11 clade, with a genome of only ∼1.3 Mbp, makes up 25% of all planktonic bacteria [24]. Streamlining is also prevalent in soils, as represented by the recently described *Candidatus Udaeobacter copiosus*, a ubiquitous taxon in soils, with a genome size of 2.81 Mbp [25].

Temperature can also influence genome size due to increased fitness of small cells at high temperatures [14]. Accordingly, small cells and smaller genomes are typically associated with higher optimal growth temperatures. This relationship is most pronounced in thermophilic communities [26], but has also been demonstrated in marine systems [27–29] and more recently in soils [30]. These patterns between genome size, GC content, and number of 16S rRNA gene copies as a result of temperature-induced genome reduction often resemble patterns in streamlined genomes [14].

Small genomes are also prevalent in host-associated bacteria, however the processes underpinning the reduction in genome size involve several mechanisms, including drift, rapid mutation rate, or other mechanisms, which could be more important than streamlining [9]. In environments where nutrients are abundant but population sizes small, deletions in bacterial genomes are more likely to become fixed in a population [9, 31], a process particularly common in host-associated gut microbiota, where population sizes are small due to isolation [32]. Bacteria subject to higher levels of mutation are more likely to be AT-rich since there is a mutational bias from GC → AT [9, 33–35]. Since the mechanisms driving the evolution of host-associated bacteria often stray from streamlining, genome reduction in host-associated bacteria may yield different patterns in genome reduction. Specifically, streamlining, which is more a directional rather than stochastic process, will often select for specific genes [9].

Much of our knowledge concerning bacterial genomic traits has been derived from cultures or isolates. This present us with substantial bias in our understanding of these relationships [36], especially for genomic traits of bacteria in complex microbial communities [37]. An alternative approach is to examine genome size on a community level *in situ*. While metagenome-assembled genomes can provide astounding insights into evolutionary processes [38], assembly can be difficult and computationally expensive, making this approach difficult to apply across a large number of communities. Fortunately, genomic traits like GC content, and average genome size, often can be easily estimated, and do not require an extensive knowledge of the taxa within the community. Here we present a comparison of genomic traits from 118 metagenomes from soil, marine, host-associated, and thermophilic systems. We hypothesize that the average genome size in soil microbial communities will be larger than in marine, host-associated, or thermophilic communities, consistent with findings from isolates [14]. We also predict that average genome size and GC content will be positively correlated in free-living soil, marine, and thermophilic communities, consistent with streamlining. Finally, we predict that while both free-living and host-associated communities with small average genome sizes will demonstrate a low GC content, free-living communities will also exhibit additional streamlined traits such as a reduced number of σ-factor and rRNA gene copies.

## METHODS

### Dataset Curation

Metagenomes from soil, marine, thermophilic, and host-associated communities were downloaded from the Integrated Microbial Genomes & Microbiomes (IMG/M) [39] system. We searched for soil and marine samples that were untreated and from natural systems (i.e. not an incubation or microcosm). For thermophilic samples we searched for communities derived from natural hot-springs, and for host-associated samples we included communities associated with an animal symbiont. We then selected samples which were both sequenced and assembled by the Joint Genome Institute (JGI) and where > 35 Mbp were assembled. Replicates appearing to be derived from a single sample (i.e. identical metadata and sample name) were discarded. In order to limit potential bias introduced by a specific study site or set of protocols of a given study, no more than 4 samples were used from any single geographical location and no more than 14 samples were selected from a single study. Ecosystem type was determined for soil samples using the available metadata and study description. Samples from non-published metagenomes were included with the consent of the primary investigator of the study. In total, 116 samples from 30 different studies were used in this analysis (Supplemental Fig. 1; Supplemental Table 1; [15, 40-62]).

### Collection of Genomic Traits

Average genome size for each metagenome was estimated using the program MicrobeCensus (parameters -n 50000000) [63] on QC filtered reads accessed through the JGI Genome Portal [64]. MicrobeCensus uses the abundance of single-copy genes to estimate the number of individuals in a population, which is then divided by the total number of read base-pairs to provide an estimate of the average genome size in a metagenome.

From IMG/M, we accessed the size of the metagenomic sample (bp), GC-%, total number of 16S rRNA gene copies, and the total number of σ factors identified by the KEGG Orthology database (Table 1 [65]). We estimated the number of genomes per metagenome by dividing the total base pair count of the metagenome by the estimated average genome size from MicrobeCensus. The average number of 16S rRNA gene copies per genome and the number of σ-factors gene copies per genome was then determined by dividing the total number of 16S rRNA or σ-factor gene copies by the estimated number of genomes.

**Table 1:**
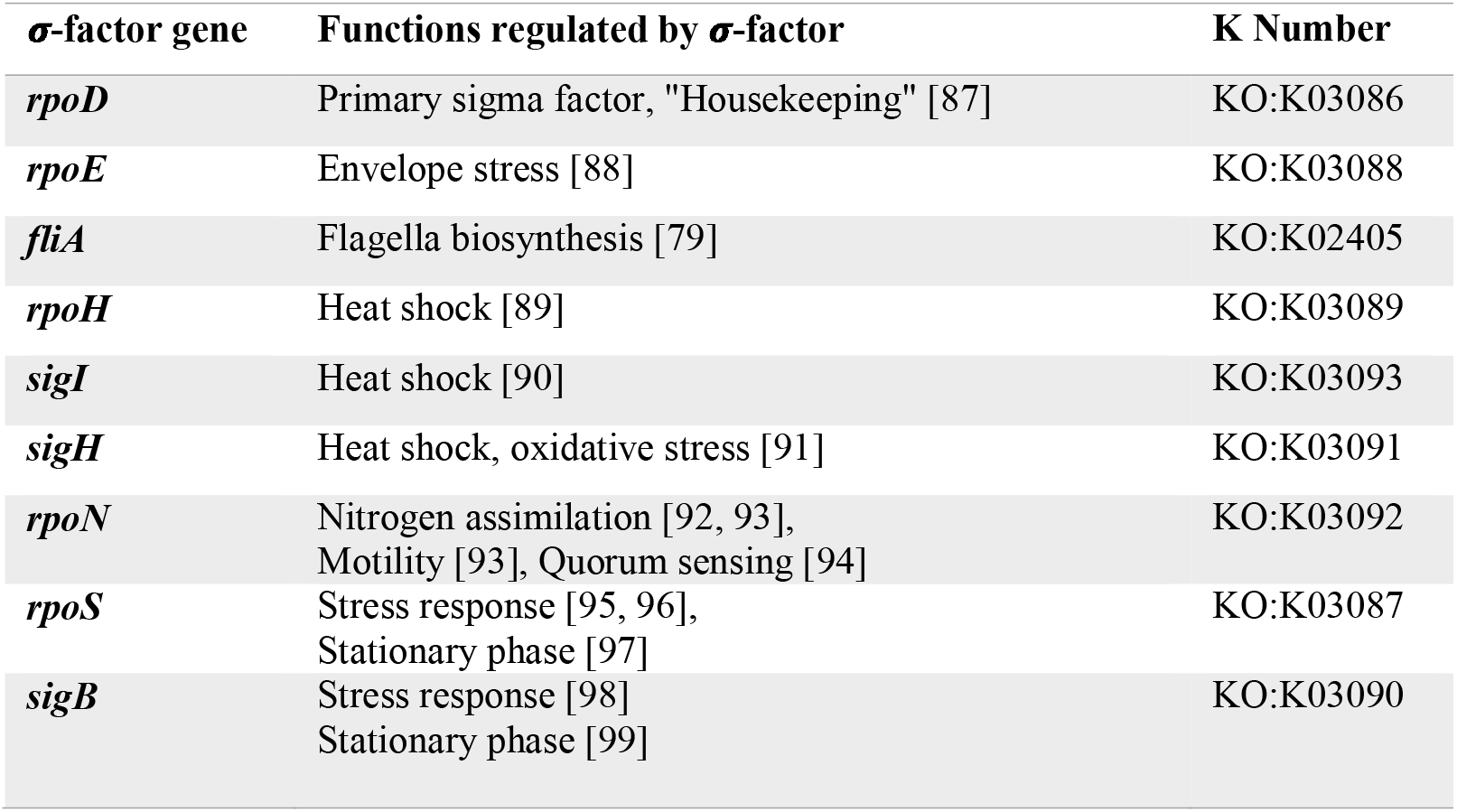
Gene name, description, and KEGG ortholog identifier (K numbers) for each σ-factor used in the analysis.

### Data Filtering

To ensure that any observed trends were not heavily influenced by the abundance of nonbacterial genomes, such as large eukaryotic genomes, we assessed the relationship between average genome size and the relative abundance of assembled bacterial reads. The phylogenetic distribution of assigned reads for each metagenome were downloaded from IMG/M and grouped by domain. The relationship between the relative abundance of bacteria and average genome size of the community was then calculated for each ecosystem to assign a cutoff which demonstrated the least amount of bias (as determined by linear regression). As a result, samples where bacteria made up less than < 95% of the assembled reads were discarded.

Since archaeal abundance in thermophilic microbial communities is often high, filtering samples with < 95% bacterial reads discarded a large number of thermophilic samples. Post filtering, only 5 thermophilic samples were left for analysis – a sample size ultimately too small to generate conclusions. Rather than omitting the thermophilic environments from our analysis entirely, and because small archaeal genomes abundance have been shown to be correlated with higher optimum growth temperatures [14], we decided to include thermophilic samples with > 5 % archaeal abundance in several of the comparisons. Although these data do not examine bacterial streamlining specifically, we find that they still provide valuable insight into how genomic traits are distributed in these communities. Mixed thermophilic samples (those including > 5 % archaea) are shown separately in figures and analyses. In comparisons of genome size versus bacteria-specific traits, such as 16S rRNA gene copies or abundance of sigma factors, we only report samples where bacteria comprise > 95% of annotated reads.

### Analysis

Multiple regression was used to determine the relationship between genome size and genomic characteristics – specifically, GC content, 16S rRNA gene relative abundance, the relative abundance of σ-factor genes, and the relative abundance of specific σ-factor variants. Models were constructed with the command lm or lmer from the R (v3.6.1 [66]) package lme4 [67]. For each response variable, we constructed multiple models considering all parameters and interactions. Final models were selected using Akaike information criterion (AIC) values. The addition of a new parameter resulting in a reduction of the AIC value by at least 4 indicated a significantly better fit with increased model complexity.

To assess the abundance of σ-factor genes between different ecosystems, we used both the multi-response permutation procedure (MRPP) as well as the permutational multivariate analysis of variance (PERMANOVA). The MRPP was conducted using all samples while PERMANOVA was conducted using 11 randomly selected genomes from each ecosystem to ensure balanced design. Both analyses were conducted using Bray-Curtis dissimilarity matrices constructed from the relative abundance of each σ-factor. To visualize differences in the distribution of different types σ-factors between ecosystems we used nonmetric multidimensional scaling (NMDS) on Bray-Curtis distances. MRPP, PERMANOVA and NMDS were done using the *vegan* package [68] in R (v3.6.1).

### Isolates

To compare relationships between genomic characteristics of a microbial community with characteristics of isolates, we accessed over 6,000 isolates of bacteria, archaea, and fungi from the IMG/M system in June of 2020. Isolates were selected if they were (1) publicly available; (2) previously published; (3) sequenced by JGI. Metadata was used to group samples into one of three ecosystem types: soil, marine, thermophilic, or host-associated. To avoid potential bias introduced by large studies selecting for specific taxa, we randomly selected no more than 20 isolates from a single study. Relationships between genomic characteristics were analyzed using multiple regression analyses as described above for the analysis of community-level traits. ANOVA was used to assess differences in the distribution of genomic characteristics between isolates and metagenomic averages.

## RESULTS

### Average Genome Size and GC Content

Average genome size was significantly different between ecosystems (ANOVA; F_4,111_ = 135.9, p < 0.01). Specifically, average genome size was higher in soils compared to marine, host-associated, or thermophilic communities (Fig. 1a, Tukey’s HSD p < 0.01). GC content (%) varied between each ecosystem (ANOVA; F_4,111_ = 140.3, p < 0.01), and was highest in soil, followed by thermophilic, host-associated, and then marine communities (Fig. 1b). Overall, GC content and average genome size were positively correlated; however, this trend varied between ecosystems (Fig. 1c). A comparison of multiple models, using AIC values as selection criteria, indicated that GC content was best predicted by average genome size, ecosystem, and their interaction (F_9,106_ = 136.1, p < 0.01, Supplemental Table 2). Specifically, GC content was positively correlated with average genome size in marine and thermophilic communities, negatively correlated in soil communities, and not significantly related in host-associated communities (Fig. 1c). The relationship between average genome size and GC content was offset between marine and thermophilic communities: thermophilic communities had a higher GC content than marine communities with the same average genome size (Fig. 1c). The relationship between GC content and average genome size was strongly driven by the abundance of archaea in the mixed thermophilic samples (Supplemental Fig. 2). In soils, average genome size and GC content were significantly different between ecosystem types (ex. Deserts, grasslands, forests; ANOVA: Mbp - F_7,38_ = 24.35, p < 0.01; GC-% - F_7,38_ = 4.986, p < 0.01; Fig. 2).

**Figure 1:**
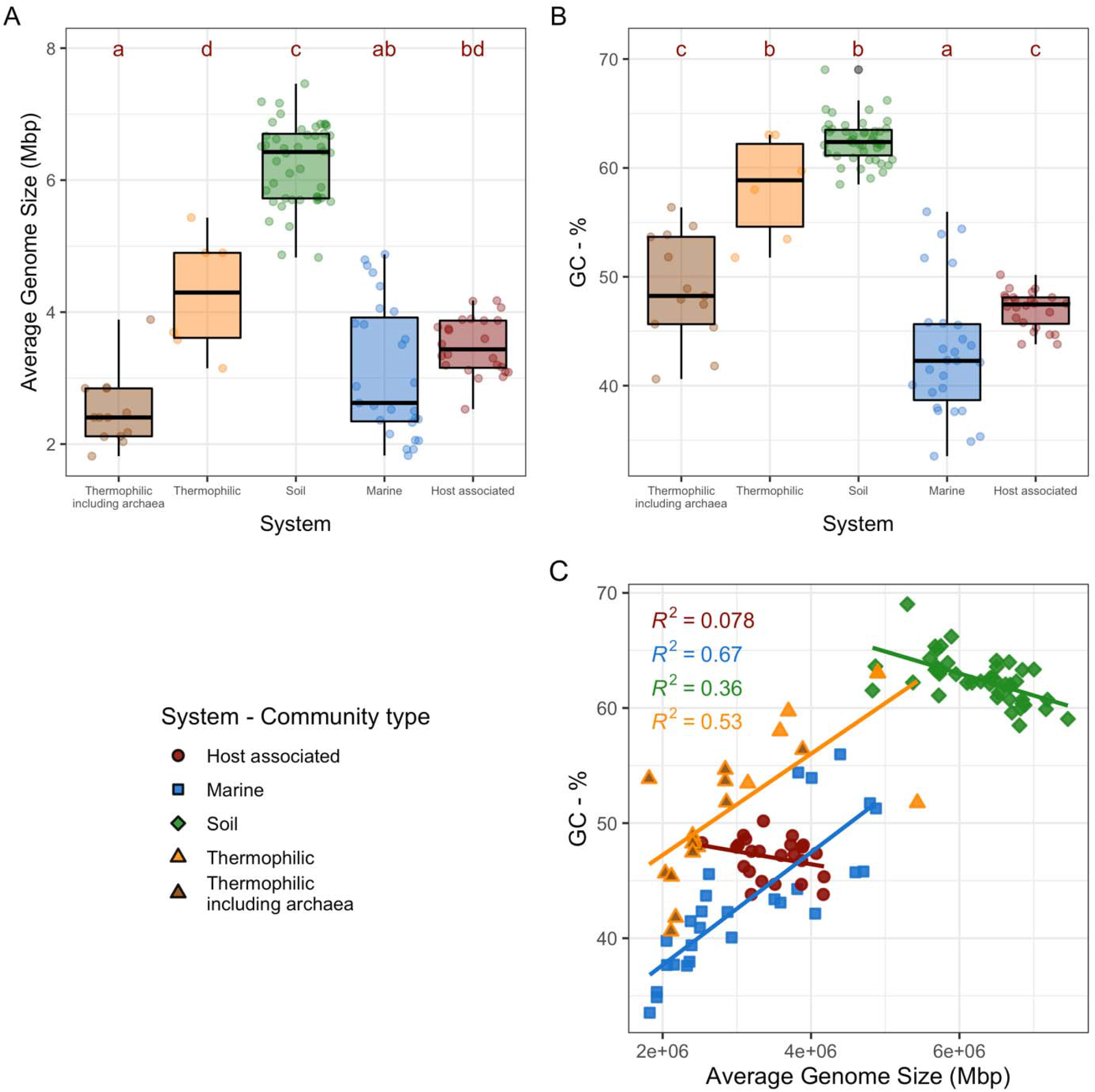
Average genome size and GC-content calculated from environmental metagenomes. (**A**) Violin-plots and boxplots of the average genome size (Mbp) of microbial communities in different ecosystems. (**B**) Violin-plots and boxplots showing GC-% between systems. **(C)** GC-% as a function of average genome size (Mbp) of a metagenome, separated by system. Point shape and outline represent source system; point fill represents system including thermophilic samples with archaea.

**Figure 2:**
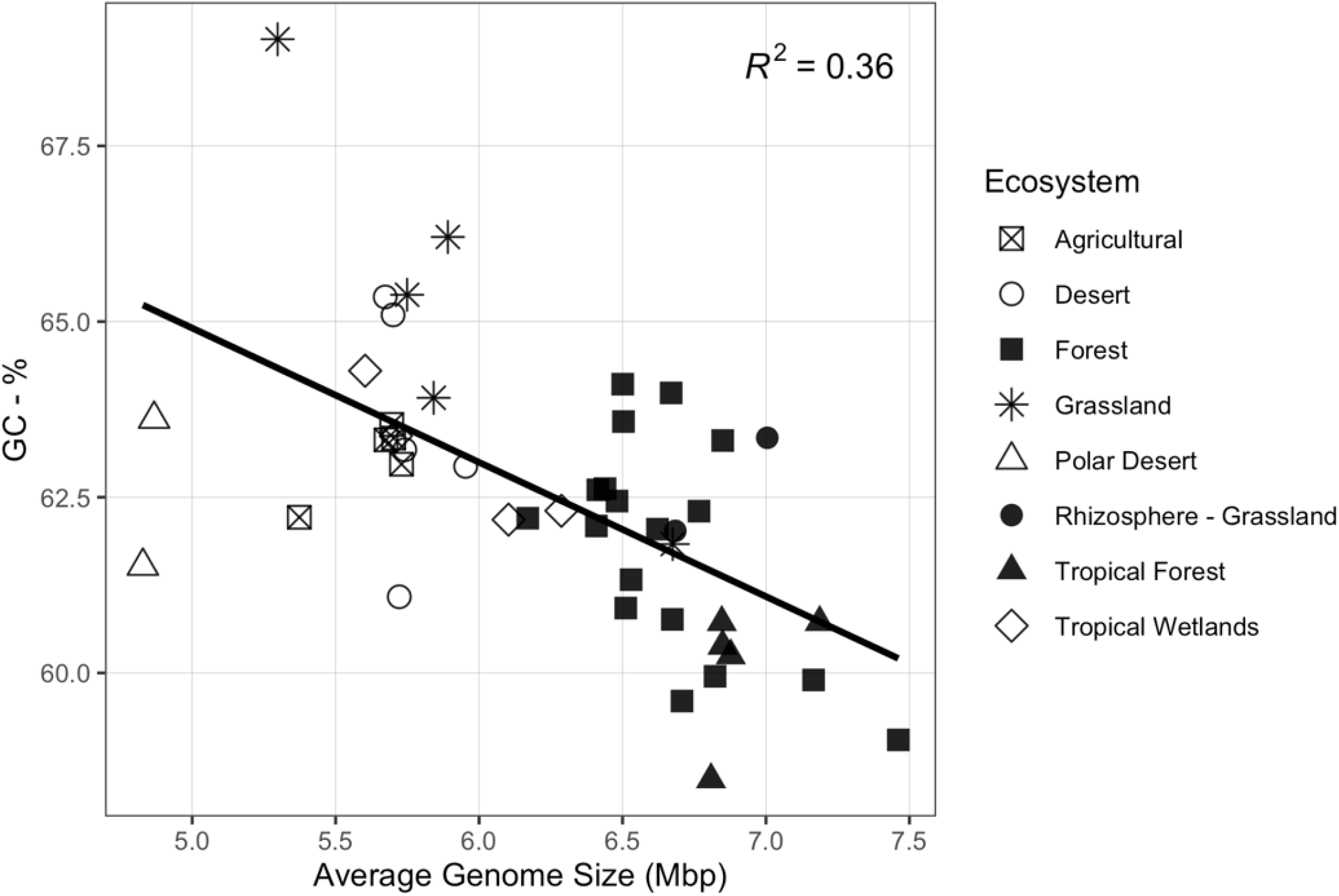
GC content (%) as a function of average genome size (Mbp) in soils, with color indicating source environment.

The average genome size and GC content of the metagenomes fell within the range of isolates from each ecosystem (Supplemental Fig. 3). However, the mean genome size and GC content derived from metagenomes varied from isolates in both soil and thermophilic environments (ANOVA; p < 0.05), but not in marine environments.

### 16S rRNA gene copies and Sigma factors

Host-associated communities had the highest number of 16S rRNA gene copies per genome, followed by soils and then thermophilic and marine communities (Supplemental Fig. 4). A comparison of AIC values indicated that ecosystem type alone was the best predictor of 16S rRNA gene copies per genome (Supplemental Fig. 4, Supplemental Table 2).

The relative abundance of σ-factors genes per metagenome changed with estimates of average genome size, however this relationship varied significantly between ecosystems (Fig. 4a; Supplemental Table 2). Average genome size was significantly correlated with the relative abundance of σ-factors in thermophilic environments (R^2^ = 0.49), but not in soil, marine, or host associated environments (R^2^ < 0.2; Fig. 4a). The distribution of σ-factor types within a metagenome varied more between ecosystems than within (Fig 3; Fig. 4b; MMRP, A = 0.34, p < 0.01), and ecosystems differed significantly (Fig 3; Fig. 4b; PERMANOVA, R^2^ = 0.50, p < 0.01).

**Figure 3:**
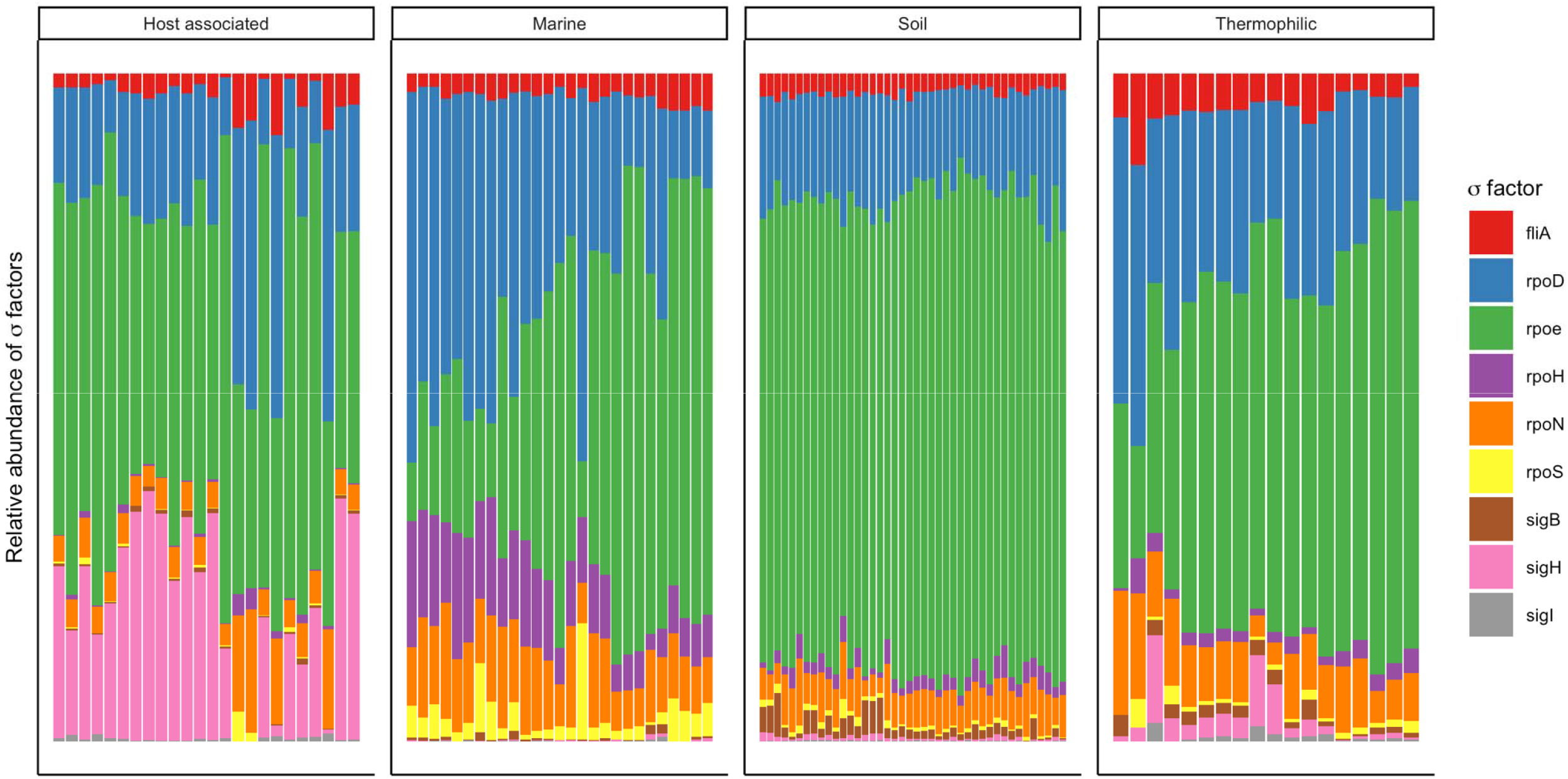
The relative abundance σ-factors in a metagenome separated by ecosystem. Each bar represents the abundance of σ-factors in a single metagenome, and metagenomes are ordered from smallest to largest (left to right) for each ecosystem.

**Figure 4:**
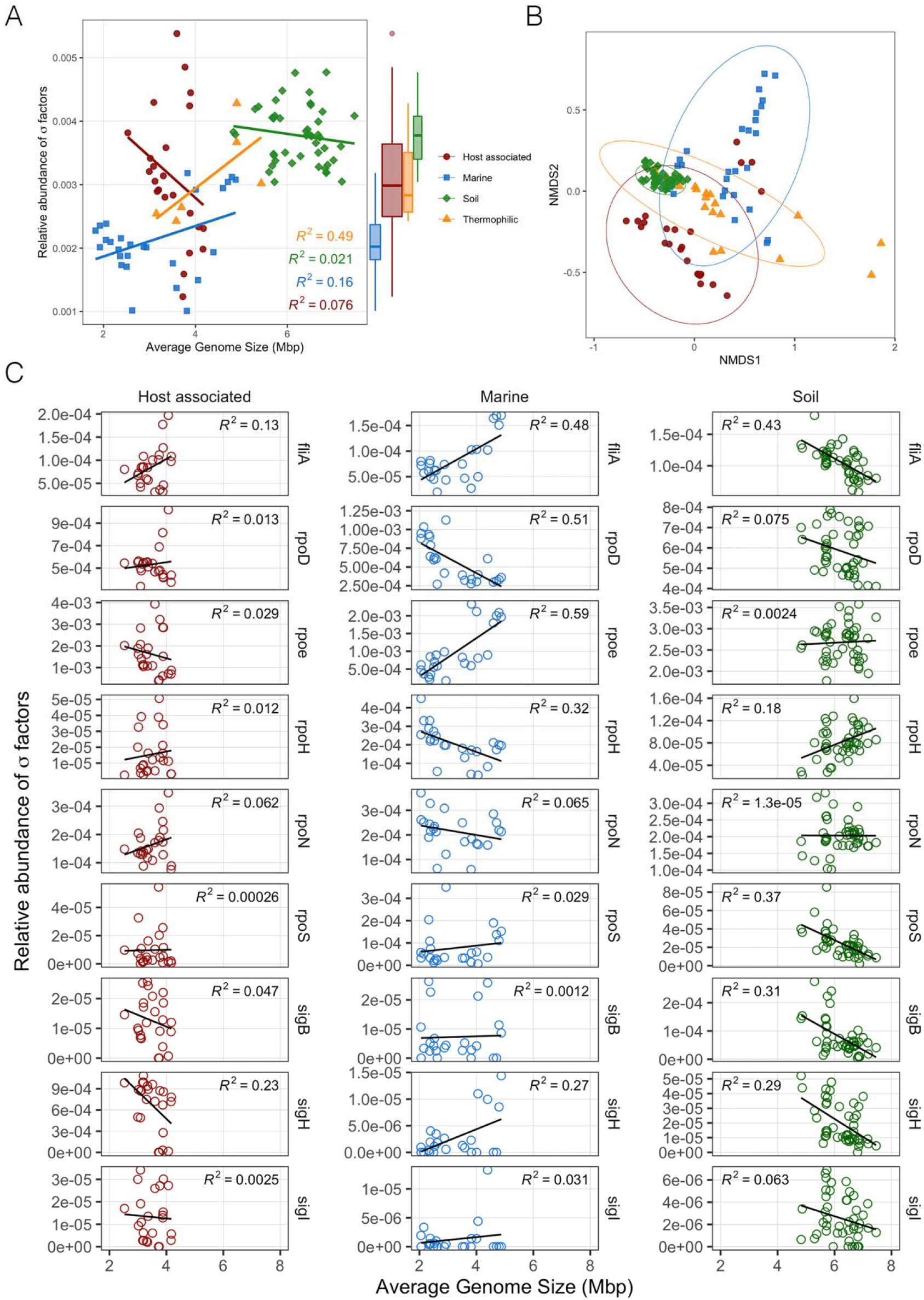
The relative abundance of σ-factors (σ-factor count / gene count) as a function of average genome size and system. **(A)** The relative abundance of all σ-factors (total σ-factor count / gene count) in a metagenome against average genome size. Source environment indicated by color for host associated (red), soil (green), thermophilic (orange) and marine (blue) communities. **(B)** NMDS of Bray-Curtis distance of the relative abundance of σ-factors (σ-factor count / total gene count) from a metagenome. **(C)** The relative abundance (σ-factor count / total gene count) of 9 σ-factors (rows) versus average genome size, separated by environment (columns).

The relationship between average genome size and the relative abundance of individual σ-factors was dependent on both ecosystem type and the type of σ-factor (Fig. 4c, Supplemental Table 3). In host-associated communities, the relative abundance of only one σ-factor, *sigH*, was significantly (P = 0.018) negative correlated with average genome size. Abundance of all other sigma factors were unchanged with genome size in host-associated communities (Supplemental Table 3). In soil communities the relative abundance of *rpoH* per metagenome significantly increased (P < 0.01) with larger average genome size, while the relative abundance per metagenome of *rpoS, sigH, sigB*, and *fliA* decreased (P < 0.01). In marine communities, we found that the relative abundance of *fliA, rpoE*, and *sigH* significantly increased (P < 0.01) with genome size, and the abundance of *rpoH*, and *rpoD* significantly decreased (P < 0.01). Due to the small samples size of thermophilic communities, we did not include the relationships between σ-factors and average genome size for thermophilic environments; however, correlation coefficients and statistics for all linear regressions between average genome size and σ-factor abundance for each ecosystem can be found in Supplemental Table 3. A visualization of average σ-factor copies per genome can be found in Supplemental Fig. 5.

## DISCUSSION

Our hypotheses were that: 1) average genome size in soil microbial communities would be higher than in marine, thermophilic, and host-associated microbial communities; 2) GC content would be positively correlated with the average genome size of free-living (not host-associated) microbial communities; and 3) host-associated microbial communities exhibiting smaller average genome sizes, although AT-rich, will lack other traits associated with streamlining, such as a reduction in regulatory σ-factors. The first and third hypotheses were generally supported, however the second hypothesis was rejected since trends between genomic traits varied between environments.

As expected, microbial communities in marine, host, and thermophilic environments had a smaller average genome size than those in soil, consistent with previous findings from studies using bacterial isolates and single-amplified genomes [8, 11, 69]. Since smaller genomes tend to have lower GC content [70], we expected to find a positive correlation between GC content and average genome size for each ecosystem. Surprisingly we only found this relationship in marine and thermophilic communities. In thermophilic communities, this relationship appeared confounded with the presence of archaea, thus making it impossible to distinguish between archaeal abundance or streamlining as a driver for smaller genome size in these extreme environments. It is worth noting that the relationship between genome size and GC content in thermophilic communities was offset higher from marine systems, even for bacterial dominated thermophilic communities. This offset is likely the result of a requirement for thermal stability in hot environments which is provided by the GC triple-hydrogen bonds versus the AT double-bond [71, 72].

Both GC content and average genome size in host-associated communities were low, a common feature of symbiotic bacteria [32]. Although host-associated bacteria in small populations often have AT-rich genomes [9], the relationship between GC content and average genome size was not significant for host-associated communities. Reduced genetic flow in these communities could mean that changes in nucleotide frequency and genome size develop independently in populations. Therefore, these trends might exist within, but not between, communities. In other words, host-associated environments might produce small AT-rich genomes, but these two traits do not covary between communities.

Soil communities exhibited a negative relationship between average genome size and GC content. This does not necessarily exclude streamlining as a driver of genome size in soils but suggests other drivers of genome size and GC content. For instance, fungal reads may reduce the overall GC content of a metagenome while raising estimates of average genome size. We found that fungal isolates generally have a lower GC content than bacteria with a similarly sized genome (Supplemental Fig. 6). Although we attempted to avoid the influence of fungal genomes by limiting our dataset to metagenomes which were dominated by bacteria, it is possible that even a low abundance of large fungal genomes affected our estimates.

Another explanation is that soil microbial communities in nutrient limited environments select for smaller genomes with a higher GC content, which may be advantageous when carbon is limited [73]. A GC basepair has carbon to nitrogen ratio of 9:8 while an AT basepair has a ratio of 10:7. A reduction in GC content therefore decreases the amount nitrogen required for DNA synthesis, which has been suggested as an explanation of the low GC content that is commonly exhibited in marine systems, where nitrogen is limiting [74]. Similarly, small genomes in soil, where C is generally limiting [75, 76], might preferentially select for GC rich DNA. In this dataset, communities from deserts, agricultural fields, and grasslands had a smaller average genome size and higher GC content (Fig. 2). These environments tend to have lower soil and microbial carbon to nitrogen ratios than forests [77]. Similarly, bacterial communities in forests tended to have larger average genome sizes and lower GC content. Other environmental factors fall along this gradient which might also influence GC content, such as temperature and moisture, which has been shown to influence nucleotide composition in terrestrial plants [78]. Our work shows a potential connection between nucleotide frequency and ecosystem properties, emphasizing the need to develop a more complete understanding of genomic features across soil microbial communities.

We did not find that the relative abundance of σ-factors was associated with average genome size. However, we did observe that marine communities maintained a lower abundance of σ-factor gene copies in comparison to other ecosystems, even when average genome size was comparable. One explanation is that the reduction of σ-factor gene copies is particularly effective in reducing reproductive costs in marine systems. Marine systems are considered to be nutrient poor relative to soils and a general reduction in the proportion of σ-factors in bacterial genomes may function as an adaptation to nutrient constraints. We also found many trends between average genome size and the abundance of specific σ-factor genes in marine communities. In marine metagenomes, the relative abundance per genome of *rpoD* and *rpoH*, which encode for σ^D^ and σ^H^ respectively, was negatively correlated with average genome size. These trends are perhaps caused by the abundance of the streamlined SAR11 clade, which only contain σ^D^ and σ^H^ [24]. Conversely, the abundance of the gene *fliA*, which encodes for the σ^28^ and regulates flagella biosynthesis [79], increased with average genome size. This relationship tracks a copiotroph-oligotroph framework in marine systems, wherein nutrient scarcity selects for smaller, more streamlined, individuals while increased nutrient availability selects for larger individuals capable of chemotaxis [17, 80].

In soils, the relative abundance of many σ-factors were negatively correlated with estimates of average genome size. Most notably, we observed a decrease in the relative abundance of *rpoS* (σ^S^) but no significant change in the abundance of *rpoD* (σ^D^) with increasing average genome size. The balance between *rpoS* and *rpoD* may be a trade-off between stress tolerance and growth [81, 82]. A higher ratio of *rpoS* to *rpoD* has been shown to increase the cell’s capacity to cope with stress but limit its ability to grow on a variety of carbon sources [82– 84]. We see this reflected in the environments from which the metagenomes were samples, with microbial communities from high stress environments, such as deserts, having a higher abundance or *rpoS* compared to lower-stress carbon-rich environments, such as forests (Supplemental Fig. 7).

Surprisingly, we found a high abundance of *fliA* gene copies in soil communities with smaller genomes, several of which were sourced from desert environments. Motility may be more valuable in nutrient limited environments, whereas in environments with high nutrient inputs, nutritional competency may be more paramount. However, these results contrast with the commonly held notion that chemotaxis is most prevalent in mesic soils. One explanation is that motility may be especially important when water availability is ephemeral. A greater number of regulatory mechanisms would therefore be advantageous as it would allow for a rapid response to periodic pulses of moisture. Another possibility is that bacteria utilize biofilms surrounding fungal hyphae, or “fungal highways” [85], which could explain the persistence of flagellated bacteria even in xeric environments [86].

## Conclusion

We found a number of compelling relationships between ecosystem parameters and genomic traits of a microbial community, most notably with genome size, GC content and the distribution of σ-factors. Several of these relationships align with evolutionary mechanisms which relate to known drivers in these environments, such as streamlining in oceans and drift in endosymbionts. Still, we observed trends in soils which were not in-line with known mechanisms of genome reduction, emphasizing the need to develop an understanding of the controls of genomic features in soils. Finally, this work highlights the utility of understanding microbial communities through the lens of genomic traits in addition to genetic potential.

## DECLARATIONS

### Availability of data and materials

All data used in this analysis were accessed from the IMG/M database (https://img.jgi.doe.gov/cgi-bin/m/main.cgi). Study names and GOLD project IDs can be found in Supplementary Table 1. Data which was not both publicly available and previously published was used with permission from the listed authors on IMG/M. **This publication does not act as a primary publication for included studies and use of these data requires consent from the team which generated these data**.

### Authors’ contributions

PC gathered and analyzed the data and wrote the manuscript. PD assisted in study design, writing, and editing. BH and ES contributed to writing and editing the manuscript. All authors read and approved the final manuscript

## ACKNOWLEDGEMENTS

These sequence data were produced by the US Department of Energy Joint Genome Institute http://www.jgi.doe.gov/ in collaboration with the user community. We would like to thank the following people and projects for granting us access to their data as part of this study: Jeanette Norton, Thea Whitman, Barbara Campbell, Janet Jansson, Ramunas Stepanauskas, Thomas Bianchi, Elise Morrison, Edward DeLong, William Mohn, Jonathan Raff, Robert Kelly, Nicole Dubilier, Steve Hallam, Mak Saito, David Walsh, Roland Hatzenpichler, Brett Baker, Frank Stewart, Erik Lilleskov, Devaki Bhaya, Brian Yu, Craig Cary, New Zealand Terrestrial Antarctic Biocomplexity Survey (NZTABS) supported by Antarctica New Zealand and the University of Waikato (Hamilton, New Zealand), Rick Cavicchioli, Jim Fredrickson, Jennifer Pett-Ridge, Kelly Gravuer, Emiley Eloe-Fadrosh, Charlene Kelly, Marina Kalyuzhnaya, James Tiedje, Anthony Neumann, Andreas Brune, and Gregory Dick.

We would also like to thank Megan Foley, Anita Antoninka, Carl Roybal, and Jeff Propster for their intellectual contributions to this work.

## REFERENCES

1. Barberán A, Ramirez KS, Leff JW, Bradford MA, Wall DH, Fierer N. Why are some microbes more ubiquitous than others? Predicting the habitat breadth of soil bacteria. Ecol Lett. 2014;17:794–802. doi:10.1111/ele.12282.

2. Krause S, Le Roux X, Niklaus PA, van Bodegom PM, Lennon T. JT, Bertilsson S, et al. Trait-based approaches for understanding microbial biodiversity and ecosystem functioning. Front Microbiol. 2014;5:251.

3. Raes J, Letunic I, Yamada T, Jensen LJ, Bork P. Toward molecular traitLbased ecology through integration of biogeochemical, geographical and metagenomic data. Mol Syst Biol. 2011;7:473. doi:10.1038/msb.2011.6.

4. Fierer N, Barberán A, Laughlin DC. Seeing the forest for the genes: Using metagenomics to infer the aggregated traits of microbial communities. Front Microbiol. 2014;5:614. doi:10.3389/fmicb.2014.00614.

5. Martiny JBH, Jones SE, Lennon JT, Martiny AC. Microbiomes in light of traits: A phylogenetic perspective. Science (80-). 2015;350. doi:10.1126/science.aac9323.

6. Green JL, Bohannan BJM, Whitaker RJ. Microbial biogeography: From taxonomy to traits. Science (80-). 2008;320:1039–43. doi:10.1126/science.1153475.

7. Vieira-Silva S, Rocha EPC. The systemic imprint of growth and its uses in ecological (meta)genomics. PLoS Genet. 2010;6:1000808. doi:10.1371/journal.pgen.1000808.

8. Cobo-Simón M, Tamames J. Relating genomic characteristics to environmental preferences and ubiquity in different microbial taxa. BMC Genomics. 2017;18:499. doi:10.1186/s12864-017-3888-y.

9. Batut B, Knibbe C, Marais G, Daubin V. Reductive genome evolution at both ends of the bacterial population size spectrum. Nat Rev Microbiol. 2014;12:841–50. doi:10.1038/nrmicro3331.

10. Roller BRK, Stoddard SF, Schmidt TM. Exploiting rRNA operon copy number to investigate bacterial reproductive strategies. Nat Microbiol. 2016;1:1–8.

11. Giovannoni SJ, Cameron Thrash J, Temperton B. Implications of streamlining theory for microbial ecology. ISME J. 2014;8:1553–65. doi:10.1038/ismej.2014.60.

12. Li J, Mau RL, Dijkstra P, Koch BJ, Schwartz E, Liu X-JA, et al. Predictive genomic traits for bacterial growth in culture versus actual growth in soil. ISME J. 2019;:1. doi:10.1038/s41396-019-0422-z.

13. Schmidt R, Gravuer K, Bossange A V, Mitchell J, Scow K. Long-term use of cover crops and no-till shift soil microbial community life strategies in agricultural soil. 2018. doi:10.1371/journal.pone.0192953.

14. Sabath N, Ferrada E, Barve A, Wagner A. Growth temperature and genome size in bacteria are negatively correlated, suggesting genomic streamlining during thermal adaptation. Genome Biol Evol. 2013;5:966–77. doi:10.1093/gbe/evt050.

15. Gravuer K, Eskelinen A. Nutrient and rainfall additions shift phylogenetically estimated traits of soil microbial communities. Front Microbiol. 2017;8:1271.

16. Giovannoni SJ, Tripp HJ, Givan S, Podar M, Vergin KL, Baptista D, et al. Genetics: Genome streamlining in a cosmopolitan oceanic bacterium. Science (80-). 2005;309:1242–5.

17. Lauro FM, McDougald D, Thomas T, Williams TJ, Egan S, Rice S, et al. The genomic basis of trophic strategy in marine bacteria. Proc Natl Acad Sci U S A. 2009;106:15527–33. doi:10.1073/pnas.0903507106.

18. Karcagi I, Draskovits G, Umenhoffer K, Fekete G, Kovács K, Méhi O, et al. Indispensability of horizontally transferred genes and its impact on bacterial genome streamlining. Mol Biol Evol. 2016;33:1257–69. doi:10.1093/molbev/msw009.

19. Kurokawa M, Seno S, Matsuda H, Ying B-W. Correlation between genome reduction and bacterial growth. DNA Res. 2016;23:517–25. doi:10.1093/dnares/dsw035.

20. Klappenbach JA, Dunbar JM, Schmidt TM. rRNA operon copy number reflects ecological strategies of bacteria. Appl Environ Microbiol. 2000;66:1328–33.

21. Yooseph S, Nealson KH, Rusch DB, McCrow JP, Dupont CL, Kim M, et al. Genomic and functional adaptation in surface ocean planktonic prokaryotes. Nature. 2010;468:60–6. doi:10.1038/nature09530.

22. Kirchman DL. Growth rates of microbes in the oceans. Ann Rev Mar Sci. 2016;8:285–309. doi:10.1146/annurev-marine-122414-033938.

23. Morris RM, Rappé MS, Connon SA, Vergin KL, Siebold WA, Carlson CA, et al. SAR11 clade dominates ocean surface bacterioplankton communities. Nature. 2002;420:806–10.

24. Giovannoni SJ. SAR11 Bacteria: The Most Abundant Plankton in the Oceans. Ann Rev Mar Sci. 2017;9:231–55. doi:10.1146/annurev-marine-010814-015934.

25. Brewer TE, Handley KM, Carini P, Gilbert JA, Fierer N. Genome reduction in an abundant and ubiquitous soil bacterium ‘Candidatus Udaeobacter copiosus.’ Nat Microbiol. 2017;2:16198. doi:10.1038/nmicrobiol.2016.198.

26. Wang Q, Cen Z, Zhao J. The survival mechanisms of thermophiles at high temperatures: An angle of omics. Physiology. 2015;30:97–106. doi:10.1152/physiol.00066.2013.

27. Swan BK, Tupper B, Sczyrba A, Lauro FM, Martinez-Garcia M, Gonzalez JM, et al. Prevalent genome streamlining and latitudinal divergence of planktonic bacteria in the surface ocean. Proc Natl Acad Sci. 2013;110:11463–8. doi:10.1073/pnas.1304246110.

28. Huete-Stauffer TM, Arandia-Gorostidi N, Alonso-Sáez L, Morán XAG. Experimental warming decreases the average size and nucleic acid content of marine bacterial communities. Front Microbiol. 2016;7:730. doi:10.3389/fmicb.2016.00730.

29. Morán XAG, Alonso-Sáez L, Nogueira E, Ducklow HW, González N, López-Urrutia Á, et al. More, smaller bacteria in response to ocean’s warming? Proc R Soc B Biol Sci. 2015;282:20150371. doi:10.1098/rspb.2015.0371.

30. Sorensen JW, Dunivin TK, Tobin TC, Shade A. Ecological selection for small microbial genomes along a temperate-to-thermal soil gradient. Nat Microbiol. 2019;4:55–61. doi:10.1038/s41564-018-0276-6.

31. Mira A, Ochman H, Moran NA. Deletional bias and the evolution of bacterial genomes. Trends Genet. 2001;17:589–96.

32. McCutcheon JP, Moran NA. Extreme genome reduction in symbiotic bacteria. Nat Rev Microbiol. 2012;10:13–26. doi:10.1038/nrmicro2670.

33. Hildebrand F, Meyer A, Eyre-Walker A. Evidence of selection upon genomic GC-content in bacteria. PLoS Genet. 2010;6:e1001107. doi:10.1371/journal.pgen.1001107.

34. Hershberg R, Petrov DA. Evidence that mutation is universally biased towards AT in bacteria. PLoS Genet. 2010;6:e1001115. doi:10.1371/journal.pgen.1001115.

35. Kuo CH, Moran NA, Ochman H. The consequences of genetic drift for bacterial genome complexity. Genome Res. 2009;19:1450–4.

36. Gweon HS, Bailey MJ, Read DS. Assessment of the bimodality in the distribution of bacterial genome sizes. ISME J. 2017;11:821–4.

37. Rinke C, Schwientek P, Sczyrba A, Ivanova NN, Anderson IJ, Cheng JF, et al. Insights into the phylogeny and coding potential of microbial dark matter. Nature. 2013;499:431–7.

38. Parks DH, Rinke C, Chuvochina M, Chaumeil P-A, Woodcroft BJ, Evans PN, et al. Recovery of nearly 8,000 metagenome-assembled genomes substantially expands the tree of life. 2017. doi:10.1038/s41564-017-0012-7.

39. Chen IMA, Chu K, Palaniappan K, Pillay M, Ratner A, Huang J, et al. IMG/M v.5.0: An integrated data management and comparative analysis system for microbial genomes and microbiomes. Nucleic Acids Res. 2019;47:D666–77.

40. Ouyang Y. Agricultural nitrogen management affects microbial communities, enzyme activities, and functional genes for nitrification and nitrogen mineralization. All Grad Theses Diss. 2016. https://digitalcommons.usu.edu/etd/5068. Accessed 6 May 2020.

41. Ouyang Y, Norton JM. Short-term nitrogen fertilization affects microbial community composition and nitrogen mineralization functions in an agricultural soil. Appl Environ Microbiol. 2020;86.

42. Whitman T, Pepe-Ranney C, Enders A, Koechli C, Campbell A, Buckley DH, et al. Dynamics of microbial community composition and soil organic carbon mineralization in soil following addition of pyrogenic and fresh organic matter. ISME J. 2016;10:2918–30.

43. Maresca JA, Miller KJ, Keffer JL, Sabanayagam CR, Campbell BJ. Distribution and diversity of rhodopsinproducing microbes in the Chesapeake Bay. Appl Environ Microbiol. 2018;84.

44. Wilhelm RC, Cardenas E, Leung H, Maas K, Hartmann M, Hahn A, et al. Data Descriptor: A metagenomic survey of forest soil microbial communities more than a decade after timber harvesting Background & Summary. 2017. doi:10.1038/sdata.2017.92.

45. Cardenas E, Orellana LH, Konstantinidis KT, Mohn WW. Effects of timber harvesting on the genetic potential for carbon and nitrogen cycling in five North American forest ecozones. Sci Rep. 2018;8:1–13.

46. Wilhelm RC, Cardenas E, Maas KR, Leung H, McNeil L, Berch S, et al. Biogeography and organic matter removal shape long-term effects of timber harvesting on forest soil microbial communities. ISME J. 2017;11:2552–68.

47. Leung HTC, Maas KR, Wilhelm RC, Mohn WW. Long-term effects of timber harvesting on hemicellulolytic microbial populations in coniferous forest soils. ISME J. 2016;10:363–75.

48. Wilhelm RC, Cardenas E, Leung H, Szeitz A, Jensen LD, Mohn WW. Long-term enrichment of stress-tolerant cellulolytic soil populations following timber harvesting evidenced by multi-omic stable isotope probing. Front Microbiol. 2017;8:537.

49. Cardenas E, Kranabetter JM, Hope G, Maas KR, Hallam S, Mohn WW. Forest harvesting reduces the soil metagenomic potential for biomass decomposition. ISME J. 2015;9:2465–76.

50. Lee LL, Blumer-Schuette SE, Izquierdo JA, Zurawski J V., Loder AJ, Conway JM, et al. Genus-wide assessment of lignocellulose utilization in the extremely thermophilic genus Caldicellulosiruptor by genomic, pangenomic, and metagenomic analyses. Appl Environ Microbiol. 2018;84.

51. Hawley AK, Torres-Beltrán M, Zaikova E, Walsh DA, Mueller A, Scofield M, et al. A compendium of multi-omic sequence information from the Saanich Inlet water column. Sci Data. 2017;4:1–11.

52. Colatriano D, Tran PQ, Guéguen C, Williams WJ, Lovejoy C, Walsh DA. Genomic evidence for the degradation of terrestrial organic matter by pelagic Arctic Ocean Chloroflexi bacteria. Commun Biol. 2018;1:1–9.

53. Baker BJ, Lazar CS, Teske AP, Dick GJ. Genomic resolution of linkages in carbon, nitrogen, and sulfur cycling among widespread estuary sediment bacteria. Microbiome. 2015;3:1–12.

54. Krüger K, Chafee M, Ben Francis T, Glavina del Rio T, Becher D, Schweder T, et al. In marine Bacteroidetes the bulk of glycan degradation during algae blooms is mediated by few clades using a restricted set of genes. ISME J. 2019;13:2800–16.

55. Camargo AP, de Souza RSC, de Britto Costa P, Gerhardt IR, Dante RA, Teodoro GS, et al. Microbiomes of Velloziaceae from phosphorus-impoverished soils of the campos rupestres, a biodiversity hotspot. Sci data. 2019;6:140.

56. Beam JP, Jay ZJ, Schmid MC, Rusch DB, Romine MF, M Jennings R De, et al. Ecophysiology of an uncultivated lineage of Aigarchaeota from an oxic, hot spring filamentous “streamer” community. ISME J. 2016;10:210–24.

57. Abraham BS, Caglayan D, Carrillo N V., Chapman MC, Hagan CT, Hansen ST, et al. Shotgun metagenomic analysis of microbial communities from the Loxahatchee nature preserve in the Florida Everglades. Environ Microbiomes. 2020;15:2. doi:10.1186/s40793-019-0352-4.

58. Hervé V, Liu P, Dietrich C, Sillam-Dussès D, Stiblik P, Šobotník J, et al. Phylogenomic analysis of 589 metagenome-assembled genomes encompassing all major prokaryotic lineages from the gut of higher termites. PeerJ. 2020;2020:e8614.

59. Rossmassler K, Dietrich C, Thompson C, Mikaelyan A, Nonoh JO, Scheffrahn RH, et al. Metagenomic analysis of the microbiota in the highly compartmented hindguts of six wood-or soil-feeding higher termites. Microbiome. 2015;3:56.

60. Mushinski RM, Payne ZC, Raff JD, Craig ME, Pusede SE, Rusch DB, et al. Nitrogen cycling microbiomes are structured by plant mycorrhizal associations with consequences for nitrogen oxide fluxes in forests. 2020; October:1–15.

61. Nayfach S, Roux S, Seshadri R, Udwary D, Varghese N, Schulz F, et al. A genomic catalog of Earth’s microbiomes. Nat Biotechnol. 2020.

62. Armstrong Z, Mewis K, Liu F, Morgan-Lang C, Scofield M, Durno E, et al. Metagenomics reveals functional synergy and novel polysaccharide utilization loci in the Castor canadensis fecal microbiome. ISME J. 2018;12:2757–69.

63. Nayfach S, Pollard KS. Average genome size estimation improves comparative metagenomics and sheds light on the functional ecology of the human microbiome. Genome Biol. 2015;16:51. doi:10.1186/s13059-015-0611-7.

64. Nordberg H, Cantor M, Dusheyko S, Hua S, Poliakov A, Shabalov I, et al. The genome portal of the Department of Energy Joint Genome Institute: 2014 updates. Nucleic Acids Res. 2014;42.

65. Kanehisa M, Goto S. Yeast Biochemical Pathways. KEGG: Kyoto encyclopedia of genes and genomes. Nucleic Acids Res. 2000;28:27–30. doi:10.1093/nar/28.1.27.

66. Team RC. R: A language and environment for statistical computing. R Found Stat Comput Vienna, Austria. 2018.

67. Bates D, Maechler M, Bolker B, … SW-, 2015 U. Package “lme4.” dk.archive.ubuntu.com. 2020. http://dk.archive.ubuntu.com/pub/pub/cran/web/packages/lme4/lme4.pdf. Accessed 12 Jun 2020.

68. Oksanen AJ, Blanchet FG, Kindt R, Legen-P Minchin PR, Hara RBO, et al. vegan: Community ecology package. 2019. doi:10.4135/9781412971874.n145.

69. Raes J, Korbel JO, Lercher MJ, von Mering C, Bork P. Prediction of effective genome size in metagenomic samples. Genome Biol. 2007;8:R10. doi:10.1186/gb-2007-8-1-r10.

70. Bentley SD, Parkhill J. Comparative genomic structure of prokaryotes. Annu Rev Genet. 2004;38:771–91. doi:10.1146/annurev.genet.38.072902.094318.

71. Musto H, Naya H, Zavala A, Romero H, Alvarez-Valín F, Bernardi G. Genomic GC level, optimal growth temperature, and genome size in prokaryotes. Biochemical and Biophysical Research Communications. 2006;347:1–3.

72. Wada A, Suyama A. Local stability of DNA and RNA secondary structure and its relation to biological functions. Progress in Biophysics and Molecular Biology. 1986;47:113–57.

73. Hellweger FL, Huang Y, Luo H. Carbon limitation drives GC content evolution of a marine bacterium in an individual-based genome-scale model. ISME J. 2018;12:1180–7. doi:10.1038/s41396-017-0023-7.

74. Grzymski JJ, Dussaq AM. The significance of nitrogen cost minimization in proteomes of marine microorganisms. ISME J. 2012;6:71–80. doi:10.1038/ismej.2011.72.

75. Demoling F, Figueroa D, Bååth E. Comparison of factors limiting bacterial growth in different soils. Soil Biol Biochem. 2007;39:2485–95. doi:10.1016/J.SOILBIO.2007.05.002.

76. Hobbie JE, Hobbie EA. Microbes in nature are limited by carbon and energy: the starving-survival lifestyle in soil and consequences for estimating microbial rates. Front Microbiol. 2013;4:324. doi:10.3389/fmicb.2013.00324.

77. Xu X, Thornton PE, Post WM. A global analysis of soil microbial biomass carbon, nitrogen and phosphorus in terrestrial ecosystems. Glob Ecol Biogeogr. 2013;22:737–49.

78. Šmarda P, Bureš P, Horová L, Leitch IJ, Mucina L, Pacini E, et al. Ecological and evolutionary significance of genomic GC content diversity in monocots.Proc Natl Acad Sci U S A. 2014;111:E4096-102. doi:10.1073/pnas.1321152111.

79. Ohnishi K, Kutsukake K, Suzuki H, Iino T. Gene fliA encodes an alternative sigma factor specific for flagellar operons in Salmonella typhimurium. MGG Mol Gen Genet. 1990;221:139– 47. doi:10.1007/BF00261713.

80. Stocker R. Marine microbes see a sea of gradients. Science (80-). 2012;338:628–33. doi:10.1126/science.1208929.

81. Nyström T. Growth versus maintenance: A trade-off dictated by RNA polymerase availability and sigma factor competition? Mol Microbiol. 2004;54:855–62. doi:10.1111/j.1365-2958.2004.04342.x.

82. Ferenci T. What is driving the acquisition of mutS and rpoS polymorphisms in Escherichia coli? Trends Microbiol. 2003;11:457–61.

83. Maharjan R, Nilsson S, Sung J, Haynes K, Beardmore RE, Hurst LD, et al. The form of a trade-off determines the response to competition. Ecol Lett. 2013;16:1267–76. doi:10.1111/ele.12159.

84. King T, Ishihama A, Kori A, Ferenci T. A regulatory trade-off as a source of strain variation in the species Escherichia coli. J Bacteriol. 2004;186:5614–20. doi:10.1128/JB.186.17.5614-5620.2004.

85. Kohlmeier S, Smits THM, Ford RM, Keel C, Harms H, Wick LY. Taking the fungal highway: Mobilization of pollutant-degrading bacteria by fungi. Environ Sci Technol. 2005;39:4640–6. doi:10.1021/es047979z.

86. Pion M, Bshary R, Bindschedler S, Filippidou S, Wick LY, Job D, et al. Gains of bacterial flagellar motility in a fungal world. Appl Environ Microbiol. 2013;79:6862–7. doi:10.1128/AEM.01393-13.

87. Lonetto M, Gribskov M, Gross CA. The σ70 family: Sequence conservation and evolutionary relationships. Journal of Bacteriology. 1992;174:3843–9. doi:10.1128/jb.174.12.3843-3849.1992.

88. Hayden JD, Ades SE. The Extracytoplasmic Stress Factor, σE, Is Required to Maintain Cell Envelope Integrity in Escherichia coli. PLoS One. 2008;3:e1573. doi:10.1371/journal.pone.0001573.

89. Grossman AD, Erickson JW, Gross CA. The htpR gene product of E. coli is a sigma factor for heat-shock promoters. Cell. 1984;38:383–90.

90. Zuber U, Drzewiecki K, Hecker M. Putative sigma factor sigI (ykoZ) of Bacillus subtilis is induced by heat shock. J Bacteriol. 2001;183:1472–5. doi:10.1128/JB.183.4.1472-1475.2001.

91. Fernandes ND, Wu QL, Kong D, Puyang X, Garg S, Husson RN. A mycobacterial extracytoplasmic sigma factor involved in survival following heat shock and oxidative stress. J Bacteriol. 1999;181:4266–74. doi:10.1128/jb.181.14.4266-4274.1999.

92. Ronson CW, Nixon BT, Albright LM, Ausubel FM. Rhizobium meliloti ntrA (rpoN) gene is required for diverse metabolic functions. J Bacteriol. 1987;169:2424–31. doi:10.1128/jb.169.6.2424-2431.1987.

93. Totten PA, Cano Lara J, Lory S. The rpoN gene product of Pseudomonas aeruginosa is required for expression of diverse genes, including the flagellin gene. J Bacteriol. 1990;172:389– 96. doi:10.1128/jb.172.1.389-396.1990.

94. Heurlier K, Dénervaud V, Pessi G, Reimmann C, Haas D. Negative control of quorum sensing by RpoN (σ54) in Pseudomonas aeruginosa PAO1. J Bacteriol. 2003;185:2227–35. doi:10.1128/JB.185.7.2227-2235.2003.

95. Battesti A, Majdalani N, Gottesman S. The RpoS-mediated general stress response in Escherichia coli. Annu Rev Microbiol. 2011;65:189–213.

96. Hengge R. The general stress response in gram-negative bacteria. In: Bacterial Stress Responses. Washington, DC, USA: ASM Press; 2014. p. 251–89. doi:10.1128/9781555816841.ch15.

97. Lange R, Hengge-Aronis R. Identification of a central regulator of stationary-phase gene expression in Escherichia coli. Mol Microbiol. 1991;5:49–59. doi:10.1111/j.1365-2958.1991.tb01825.x.

98. Hecker M, Schumann W, Völker U. Heat-shock and general stress response in Bacillus subtilis. Molecular Microbiology. 1996;19:417–28.

99. Boylan SA, Redfield AR, Price CW. Transcription factor σB of Bacillus subtilis controls a large stationary-phase regulon. J Bacteriol. 1993;175:3957–63. doi:10.1128/jb.175.13.3957-3963.1993.

