## Supplemental figures and tables for "Soil, ocean, hot spring, and host-associated environments reveal unique selection pressures on genomic features of bacteria in microbial communities"

**Supplemental Table 1:**

Study name, GOLD Study ID and relevant publications of studies used in the analysis.

| GOLD Study ID | Study Name | Associated Publications |
| --- | --- | --- |
| Gs0114436 | Agricultural soil microbial communities from Utah and Georgia to study Nitrogen management | Ouyang 2016, 2020 |
| Gs0127393 | Amended soil microbial communities from New York, USA to study carbon cycling | Whitmann 2016 |
| Gs0120398 | Animal gut microbial communities from fecal samples from Wisconsin, USA |  |
| Gs0114433 | Aqueous microbial communities from the Delaware River/Bay and Chesapeake Bay under freshwater to marine salinity gradient to study organic matter cycling in a time-series | Maresca 2018 |
| Gs0111485 | Bacterial and archaeal communities from various locations to study Microbial Dark Matter (Phase II) |  |
| Gs0131207 | Coastal seawater microbial communities from Marineland, Florida, United States |  |
| Gs0128773 | Comprehensive metagenome and single cell genome sequencing from the open ocean community of North Pacfic Subtropical Gyre, Station ALOHA |  |
| Gs0084161 | Cubitermes and Nasutitermes termite gut microbial communities from Max Planck Institute for Terrestrial Microbiology, Germany | Herve 2020; Rossmassler2015 |
| Gs0071004 | Forest soil microbial communities from multiple locations in Canada and USA | Wilhelm 2017 (3); Cardenas 2015, 2018; Mohn 2017 |
| Gs0135153 | Hardwood forest soil microbial communities from various locations in the United States | Mushinski 2020 |
| Gs0095507 | Hot spring microbial communities from Yellowstone National Park | Lee 2018 |
| Gs0095504 | Marine gutless worms symbiont microbial communities from various locations |  |
| Gs0046785 | Marine microbial communities from expanding oxygen minimum zones in the northeastern subarctic Pacific Ocean |  |
| Gs0127371 | Marine microbial communities from the Costa Rica Dome and surrounding waters |  |
| Gs0110176 | Marine microbial communities from the Southern Atlantic Ocean affecting the dissolved organic carbon pool |  |
| Gs0114292 | Marine microbial communities from the Southern Atlantic ocean transect to study dissolved organic matter and carbon cycling |  |
| Gs0127566 | Methane metabolizing microbial communities from different methane-rich environments from various locations | Collection described in Armstrong 2018 |
| Gs0110190 | Microbial and viral regulation of community carbon cycling across diverse low-oxygen zones |  |
| Gs0084160 | Pelagic marine microbial communities from North Sea | Krüger 2018 |
| Gs0131210 | Plant-associated microbial communities from Velloziaceae species in rupestrian grasslands, the National Park of Serra do Cipo, Brazil | Camargo 2019 |
| Gs0053075 | Polar desert microbial communities from Antarctic Dry Valleys |  |
| Gs0114820 | Saline lake microbial communities from various lakes in Antarctica | Included in Nayfach 2020 (not primary publication of these data) |
| Gs0067861 | Saline, thermophilic phototrophic and chemotrophic mat microbial communities from various locations in USA and Mexico |  |
| Gs0120349 | Serpentine soil microbial communities from UC McLaughlin Reserve, CA, USA | Gravuer 2017 |
| Gs0136050 | Soil and marine airborne microbial communities from various locations |  |
| Gs0136122 | Soil microbial communities from Loxahatchee National Wildlife Refuge, Florida, United States | Abraham 2020 |
| Gs0133409 | Soil microbial communities from Purdue University Martell Research Forest, Indiana, United States |  |
| Gs0128792 | Systems level insights into methane cycling in arid and semi-arid ecosystems |  |
| Gs0121362 | Terrestrial soil microbial communities with and without Nitrogen fertilizer from Kellogg Biological Station, Michigan, USA |  |
| Gs0075432 | Tropical forest soil microbial communities from Luquillo Experimental Forest, Puerto Rico |  |

**Supplemental Table 2:**

Comparison of models predicting GC-%, 16s rRNA gene copy relative abundance, and sigma-factor relative abundance. Model comparisons were conducted using AIC values as selection criteria. New parameters which reduced the AIC value by > 4 were added to the model. The optimum model for each response variable is highlighted in bold.

| **Response** | **Model** | **df** | **AIC** |
| --- | --- | --- | --- |
| GC-% | System | 4 | 645.4616 |
|  | Genome Size | 3 | 701.8270 |
|  | Genome Size + System | 5 | 628.2078 |
|  | **Genome Size x System** | **11** | **570.8312** |
| Estimated 16s rRNA genes copies per genome |  |  |  |
|  | Null | 2 | 570.1399 |
|  | **System** | **5** | **518.8544** |
|  | Genome Size (Mbp) | 3 | 563.2023 |
|  | Genome Size (Mbp) + System | 6 | 516.8588 |
|  | Genome Size (Mbp) x System | 9 | 511.3246 |
| Sigma-factors (relative abundance) |  |  |  |
|  | Genome Size (Mbp) | 3 | -1183.713 |
|  | **System** | **5** | **-1206.336** |
|  | Genome Size (Mbp) + System | 6 | -1205.114 |
|  | Genome Size (Mbp) x System | 9 | -1210.136 |

**Supplemental Table 3:**

Output of linear regression models determining the relationship between genome size and the relative abundance of sigma factors for each ecosystem type. Coefficients and statistics highlighted in bold represent strongly significant relationships (P < 0.01). Bolded and italicized output represents weakly significant relationships (0.05 > P > 0.01).

|  | Host associated | | | |  | Marine | | | |
| --- | --- | --- | --- | --- | --- | --- | --- | --- | --- |
|  | B | F1,22 | R-sq | p-value |  | B | F1,25 | R-sq | p-value |
| *fliA* | 3.46E-05 | 3.26 | 0.09 | 8.48E-02 |  | **2.95E-05** | **25.14** | **0.48** | **3.60E-05** |
| *rpoD* | 3.46E-05 | 0.28 | -0.03 | 6.00E-01 |  | **-2.41E-04** | **35.32** | **0.57** | **3.33E-06** |
| *rpoE* | -3.70E-04 | 0.65 | -0.16 | 4.30E-01 |  | **5.32E-04** | **43.10** | **0.62** | **7.06E-07** |
| *rpoH* | 3.47E-06 | 0.26 | -0.03 | 6.13E-01 |  | **-7.02E-05** | **19.15** | **0.41** | **1.88E-04** |
| *rpoN* | 3.68E-05 | 1.45 | 0.02 | 2.40E-01 |  | -2.16E-05 | 2.50 | 0.05 | 1.26E-01 |
| *rpoS* | 4.77E-07 | 0.01 | -0.05 | 9.40E-01 |  | 8.83E-06 | 0.34 | -0.03 | 5.63E-01 |
| *sigB* | -3.93E-06 | 1.09 | 0.00 | 3.09E-01 |  | -8.06E-08 | 0.00 | -0.04 | 9.59E-01 |
| *sigH* | ***-0.0003879*** | ***6.39*** | ***0.19*** | ***1.92E-02*** |  | **1.89E-06** | **8.13** | **0.22** | **8.61E-03** |
| *sigI* | -1.25E-06 | 0.05 | -0.04 | 9.18E-01 |  | 5.35E-07 | 1.09 | 0.00 | 3.07E-01 |
|  | Soil | | | |  | Thermophilic | | | |
|  | B | F1,44 | R-sq | p-value |  | B | F1,4 | R-sq | p-value |
| *fliA* | **-2.51E-05** | **33.26** | **0.43** | **7.41E-07** |  | -2.66E-05 | 1.26 | 0.04 | 3.25E-01 |
| *rpoD* | -4.87E-05 | 3.57 | 0.07 | 6.55E-02 |  | -4.77E-05 | 3.74 | 0.35 | 1.25E-01 |
| *rpoE* | 3.38E-05 | 0.11 | 0.00 | 7.45E-01 |  | 6.10E-04 | 6.24 | 0.51 | 6.68E-02 |
| *rpoH* | **2.02E-05** | **9.58** | **0.18** | **3.41E-03** |  | **3.91E-05** | **52.7** | **0.91** | **1.91E-03** |
| *rpoN* | -2.77E-07 | 0.00 | 0.00 | 9.81E-01 |  | -3.22E-06 | 0.06 | -0.23 | 8.16E-01 |
| *rpoS* | **-1.44E-05** | **25.30** | **0.37** | **8.73E-06** |  | 1.05E-05 | 2.97 | 0.28 | 1.59E-01 |
| *sigB* | **-5.63E-05** | **19.36** | **0.31** | **6.79E-05** |  | -7.47E-06 | 0.33 | -0.15 | 5.29E-01 |
| *sigH* | **-1.22E-05** | **18.31** | **0.29** | **9.97E-05** |  | -1.04E-05 | 0.99 | -0.01 | 3.75E-01 |
| *sigI* | -8.15E-07 | 2.95 | 0.06 | 9.29E-02 |  | -3.01E-06 | 0.43 | -0.13 | 5.49E-01 |

Supplemental Figure 1:

Map showing the central location of each study. Study system is indicated by color and symbol shape.

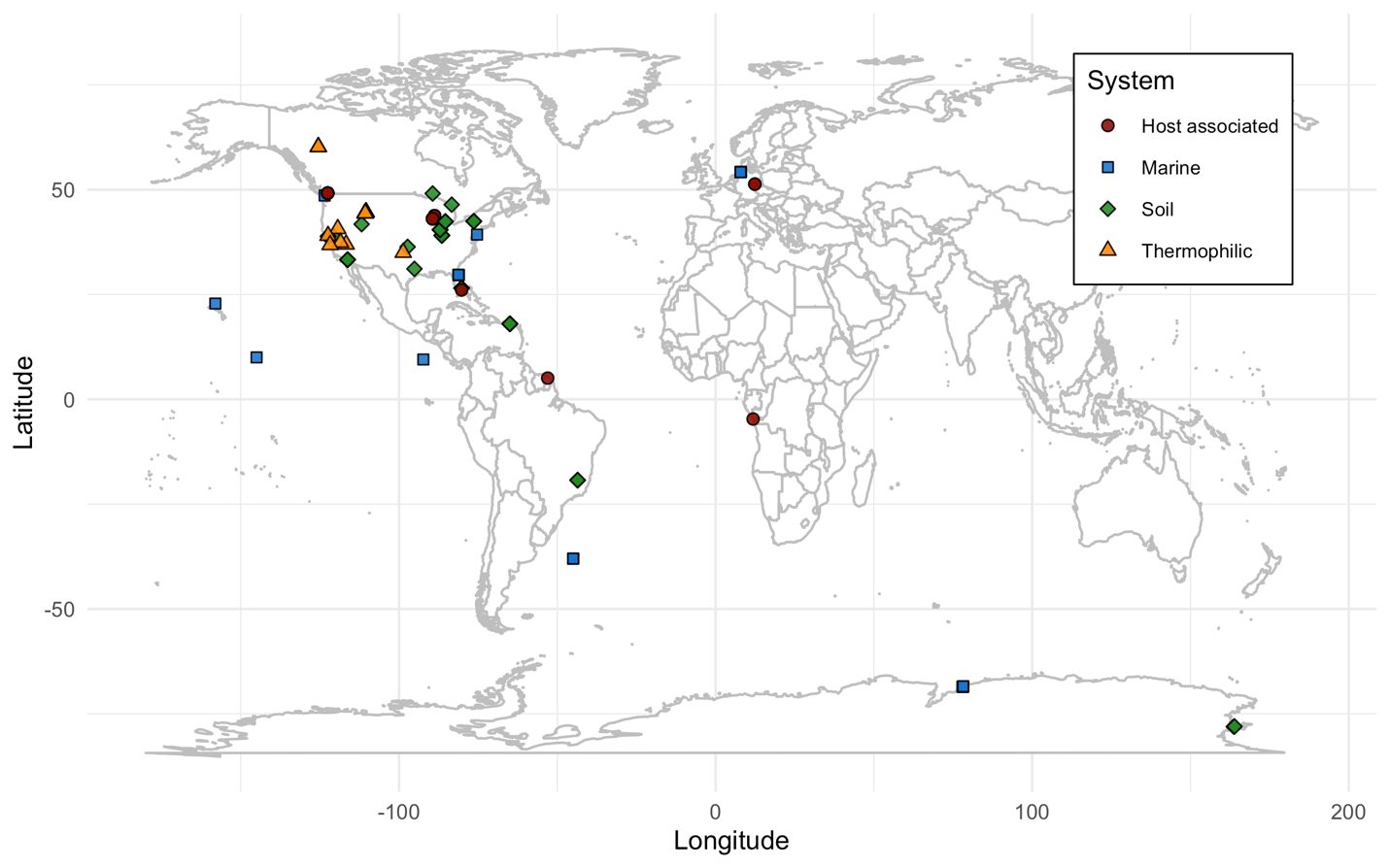

Supplemental Figure 2:

For thermophilic communities, GC content (%; **A**) and average genome size (Mbp; **B**) as a function of archaeal relative abundance (% of annotated reads).

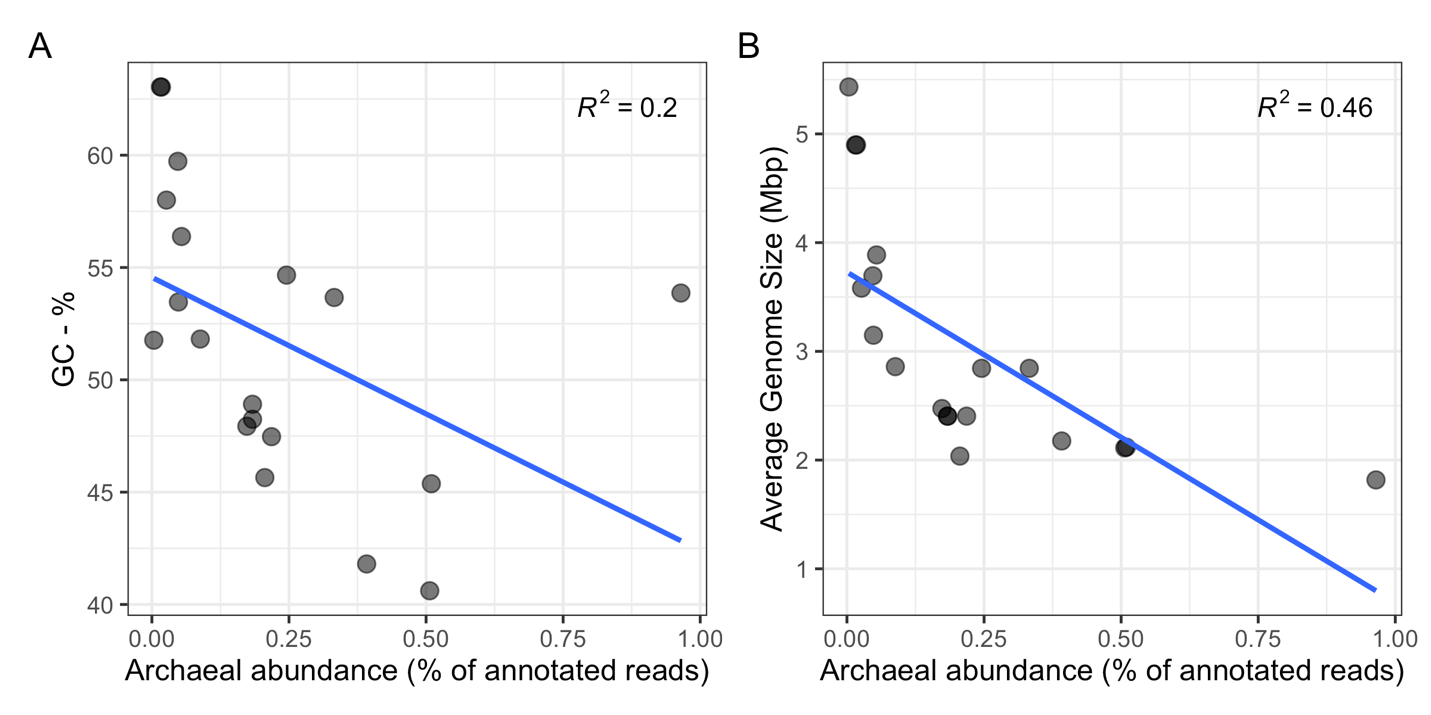

Supplemental Figure 3:

The relationship and distribution of genome size and GC content for isolates and metagenomic averages for each system. In each panel, metagenomes (dark circles) are plotted against bacterial (light squares) and archaeal (light triangles) isolates. Regression lines between genome size and GC-% are shown for both metagenomes (dark lines) and isolates (light lines). Marginal density plots show the distributions of GC-% (right) and genome size (top) for isolates (light) and metagenomic averages (dark).

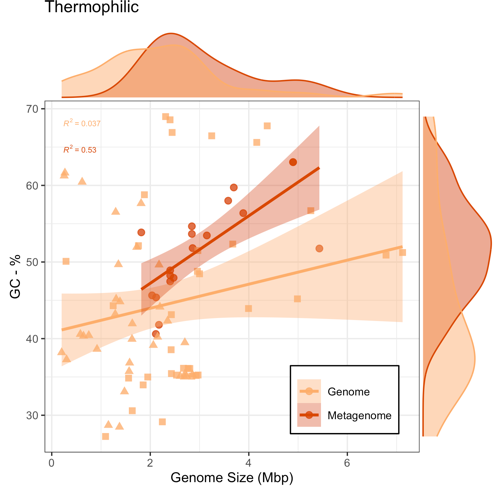

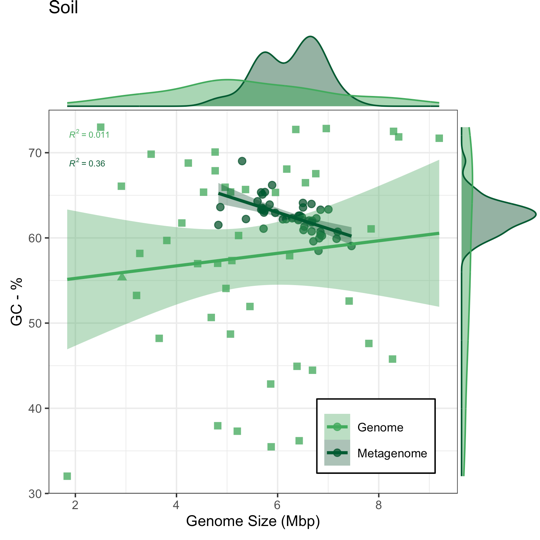

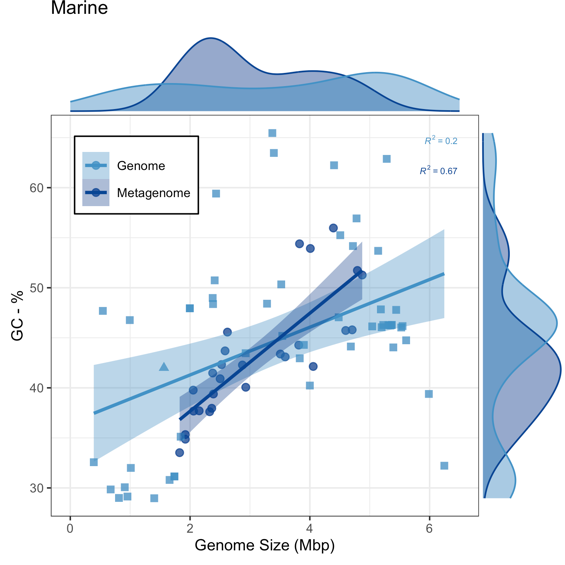

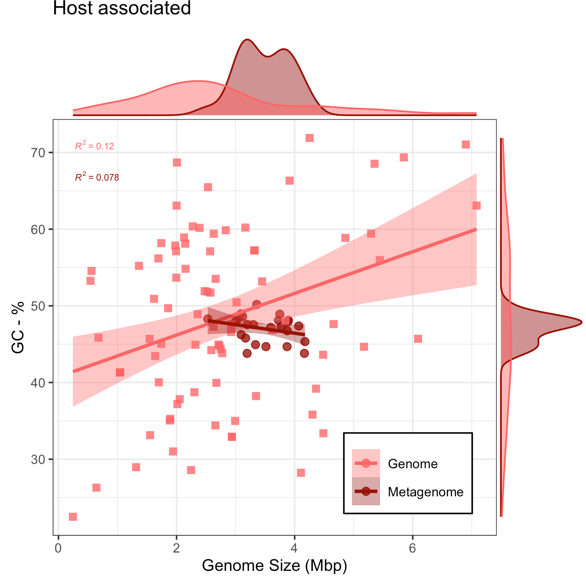

Supplemental Figure 4:

Estimated 16s copies per genome between ecosystems (**A**), and by average genome size with slopes shown separately for each ecosystem (**B**).

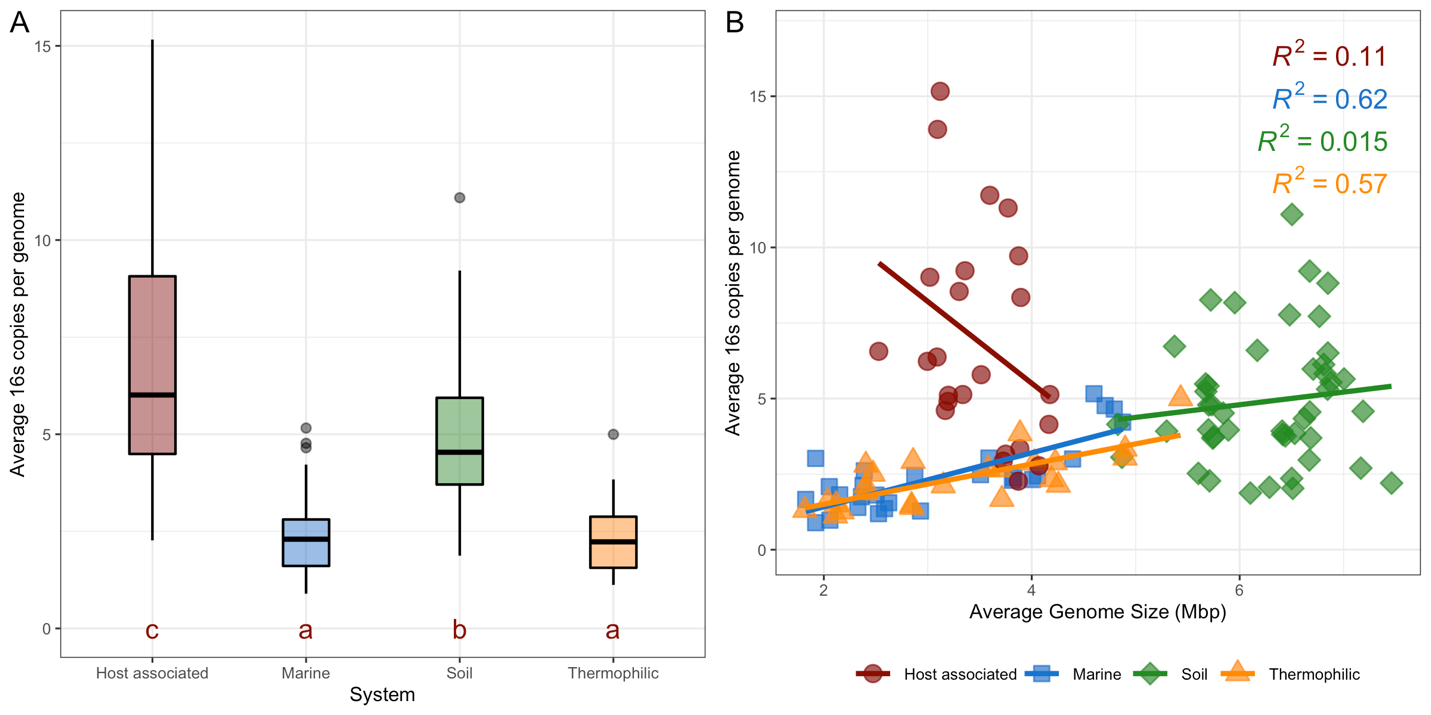

Supplemental Figure 5:

Here, and in the discussion, we discuss the relative abundance of each σ-factor as opposed to the more biologically relevant metric of average copies per genome. This is because the number of copies per genome relies on the calculation of genome size, thus resulting in autocorrelation in any analysis relating average genome size to average number of gene copies per genome. Still, we believe that the slopes for these relationships are biologically relevant and may be of interest to some readers. We therefore have provided a visualization for the average total number of σ-factors per genome (**A**), as well as the average number of specific σ-factor variants per genome (**B**).

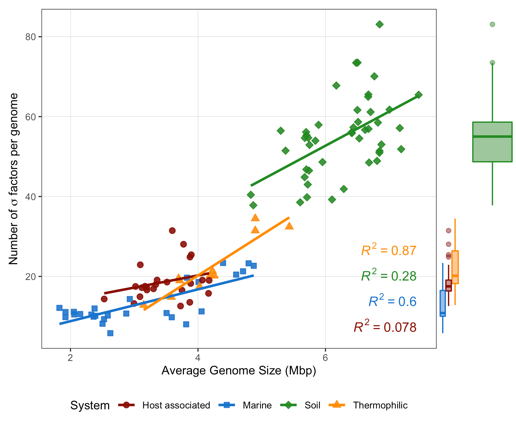

A

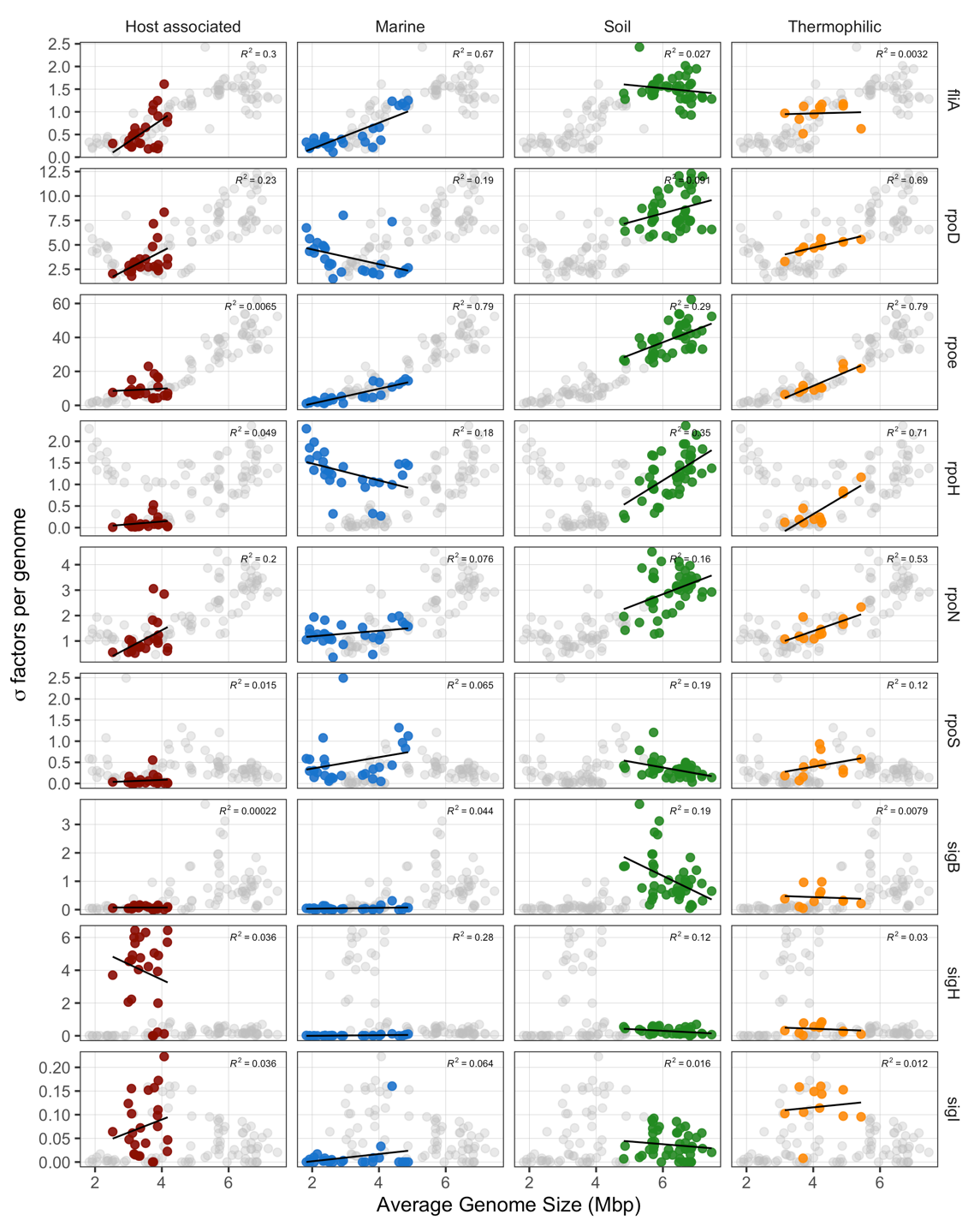

B

Supplemental Figure 6:

GC content as a function of genome size for soil archaeal, bacterial, and fungal isolates derived from the IMG database.

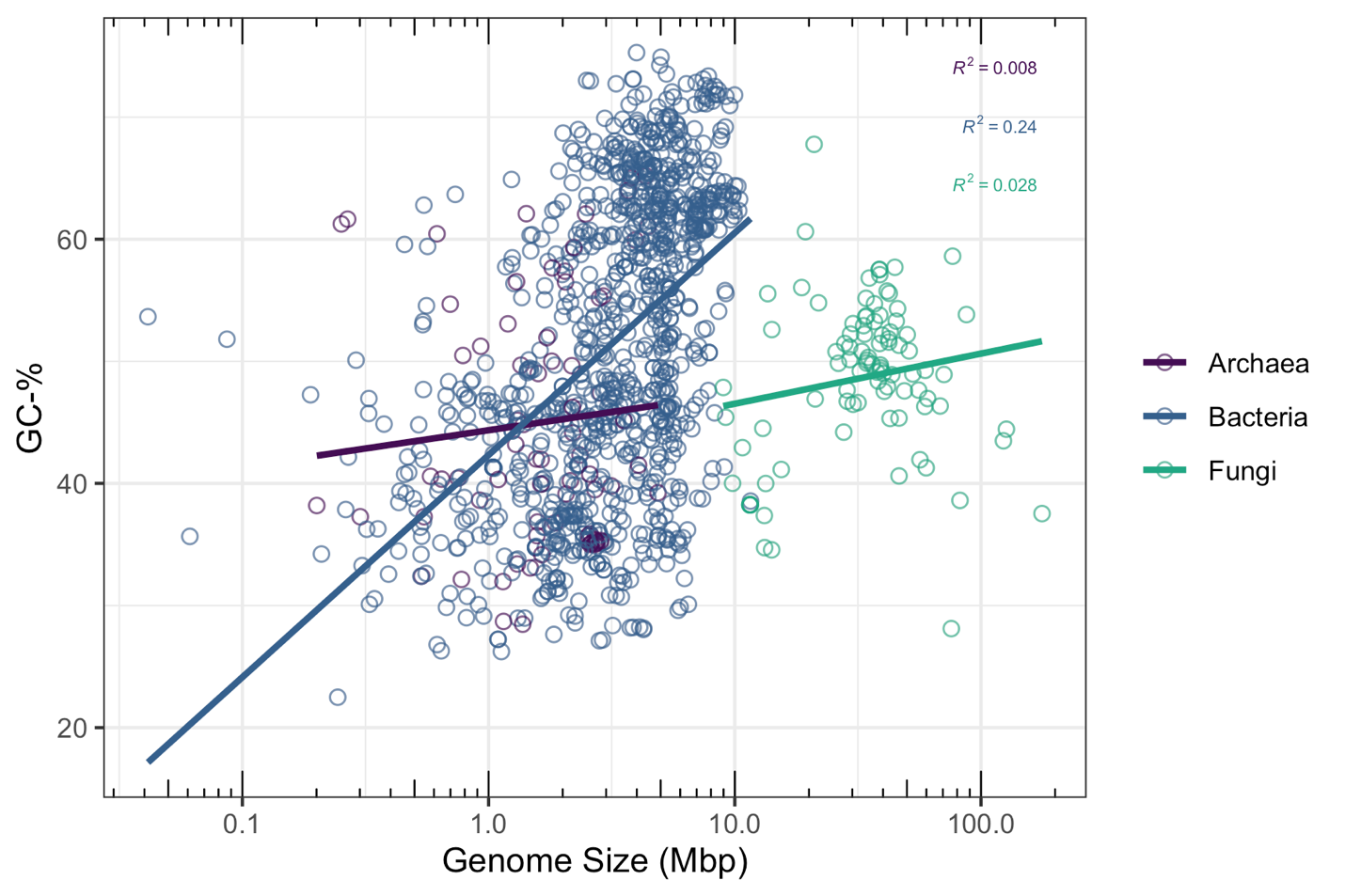

Supplemental Figure 7:

Relative abundance of σ^s^ gene *rpoS* as a function of average genome size, with source system indicated by color.

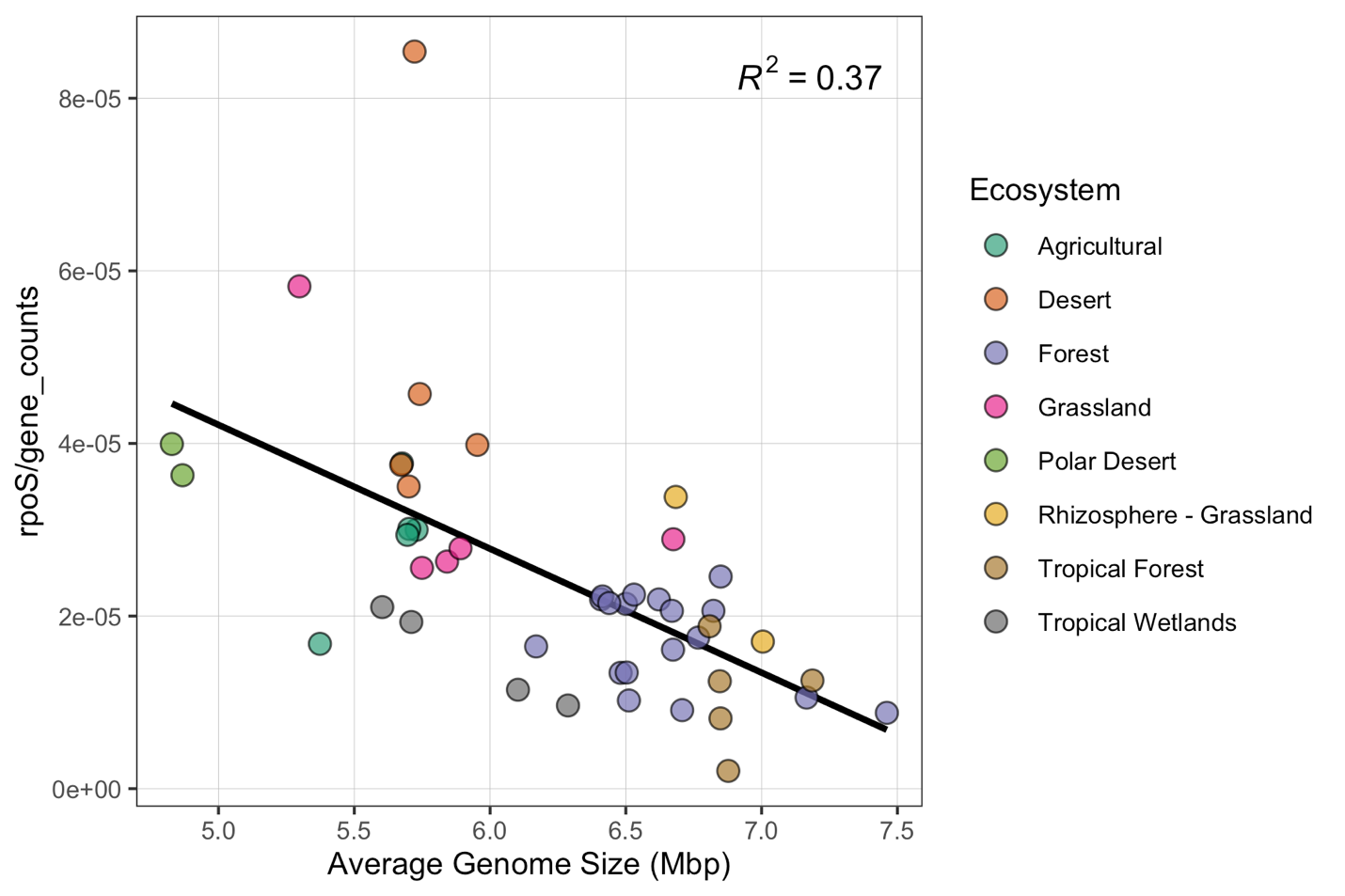
